# Genetic basis of phenotypic plasticity and genotype x environment interaction in a multi-parental population

**DOI:** 10.1101/2020.02.07.938456

**Authors:** Isidore Diouf, Laurent Derivot, Shai Koussevitzky, Yolande Carretero, Frédérique Bitton, Laurence Moreau, Mathilde Causse

**Author notes:** **Correspondence:** Mathilde Causse, INRA, UR1052, Génétique et Amélioration des Fruits et Légumes, 67 Allée des Chênes, Centre de Recherche PACA, Domaine Saint Maurice, CS60094, Montfavet, 84143, France.

## Abstract

Deciphering the genetic basis of phenotypic plasticity and genotype x environment interaction (GxE) is of primary importance for plant breeding in the context of global climate change. Tomato is a widely cultivated crop that can grow in different geographical habitats and which evinces a great capacity of expressing phenotypic plasticity. We used a multi-parental advanced generation intercross (MAGIC) tomato population to explore GxE and plasticity for multiple traits measured in a multi-environment trial (MET) design comprising optimal cultural conditions and water deficit, salinity and heat stress over 12 environments. Substantial GxE was observed for all the traits measured. Different plasticity parameters were estimated through the Finlay-Wilkinson and factorial regression models and used together with the genotypic means for quantitative trait loci (QTL) mapping analyses. Mixed linear models were further used to investigate the presence of interactive QTLs (QEI). The results highlighted a complex genetic architecture of tomato plasticity and GxE. Candidate genes that might be involved in the occurrence of GxE were proposed, paving the way for functional characterization of stress response genes in tomato and breeding for climate-adapted crop.

**Highlight:** The genetic architecture of tomato response to several abiotic stresses is deciphered. QTL for plasticity and QTL x Environment were identified in a highly recombinant MAGIC population.

## INTRODUCTION

Plants are sessile organisms which have to cope with environmental fluctuations to ensure species reproduction for persistence in nature. For a given genotype, the expression of different phenotypes according to the growing environment is commonly called phenotypic plasticity (PP) (Bradshaw, 1965). It offers the possibility to plants to adapt to new environments, notably new locations, changes in climatic conditions or seasonal variations. In agriculture, the range of environmental variation for crop cultivation may also include different cultural practices or growing conditions, leading to the expression of PP on agronomic traits and unstable performance. When different genotypes/accessions are examined for PP within a species, inter-individual variations in their responses usually lead to the common phenomenon of genotype-environment (GxE) interaction (El-Soda et al., 2014). Understanding the genetic mechanisms driving PP and GxE in plants is a crucial step for being able to predict yield performance of crop cultivars and to adapt breeding strategies according to the targeted environments.

In plants, the genetic basis of PP has been investigated to assess whether PP has its own genetic regulation and thus could be directly selected. Three main genetic models, widely known as the over-dominance, allelic-sensitivity and gene-regulatory models were proposed in the literature as underlying plant PP (Scheiner, 1993; Via et al., 1995). The over-dominance model suggests that PP is negatively correlated to the number of heterozygous loci (Gillespie and Turelli, 1989). The heterozygous status is favored by allele’s complementarity in this case. Allelic-sensitivity and gene-regulatory models are assumed to arise from the differential expression of an allele according to the environment and epistatic interactions between structural and regulatory alleles, respectively. The latter assumes an independent genetic control of mean phenotype and plasticity of a trait. Using a wide range of environmental conditions, the prevalence of the allelic-sensitivity or gene-regulatory model in explaining the genetic architecture of PP was explored in different crop species including barley (Lacaze et al. 2009), maize (Gage et al., 2017; Kusmec et al., 2017), soybean (Xavier et al., 2018) and sunflower (Mangin et al., 2017).

Quantification of PP is however a common question when analyzing the genetic architecture of plasticity since different parameters for PP estimation are available as reviewed by Valladares et al. (2006). At a population level, when multiple genotypes are screened in different environments, different approaches can be used to assess plasticity (Laitinen and Nikoloski, 2019). The most common of these approaches is the joint regression model (Finlay and Wilkinson, 1963) that uses the average performance of the set of tested genotypes in each environment as an index on which the individual phenotypes are regressed. This model, commonly known as the Finlay-Wilkinson regression model, allows to estimate linear (slopes) and non-linear plasticity parameters (from the residual errors) that presumably have different genetic basis (Kusmec et al., 2017). If the detailed description of the environments is available, the environmental index used in the Finlay-Wilkinson regression model can be replaced by environmental covariates such as stress indexes through factorial regression models (Malosetti et al. 2013). Thus plasticity could be estimated as the degree of sensitivity to a given stress continuum (Mangin et al., 2017).

Climate change is predicted to increase the frequency and intensity of abiotic stresses with a high and negative impact on crop yield (Zhao et al., 2017). Plants respond to abiotic stresses by altering their morphology and physiology, reallocating the energy for growth to defense against stress (Munns and Gilliham, 2015). Consequences on agronomic performances are apparent and detrimental to productivity. The most common abiotic stresses studied across species are water deficit (WD), salinity stress (SS) and high temperature stress (HT). The negative impact of these stresses on yield have been underlined for major cultivated crops; however, positive effects of WD and SS on fruit quality have been observed in fruit trees and some vegetables notably in tomato (Costa et al. 2007; Mitchell et al. 1991; Ripoll et al. 2014).

Tomato is an economically important crop and a plant model species which led to numerous studies that contributed much in understanding the genetic architecture of the crop and its response to environmental variation. However, most of the studies that addressed the genetic architecture of tomato response to environment were conducted on experimental populations exposed to two conditions (*i*.*e*. control *vs* stress). Albert et al. (2018) for example identified different WD-response quantitative trait loci (QTL) in a bi-parental population derived from a cross of large and cherry tomato accessions. Tomato heat-response QTLs were also identified in different experimental populations including interspecific and intraspecific populations (Grilli et al., 2007; Xu et al., 2017a; Driedonks et al., 2018). These studies investigated heat-response QTLs using mostly reproductive traits screened under heat stress condition. Villalta et al. (2007) and Diouf et al. (2018) investigated the genetic architecture of tomato response to SS and identified different QTLs for physiological and agronomic traits, involved in salinity tolerance. However, no QTL study has yet been conducted on tomato plasticity assessed under a multiple stress design, although the coincidence of different stresses is a more realistic scenario in crop cultivation, especially with the climate change.

Tomato benefits of a large panel of genetic resources that have been used in multiple genetic mapping analyses (Grandillo et al. 2013). Bi-parental populations were first used in QTL mapping and permitted the characterization of plenty of QTLs related to yield, disease resistance and fruit quality. In the genomic era, new experimental populations were developed offering higher power and advantages for QTL detection. These include mutant collections, BIL-populations and multi-parent advanced generation intercross (MAGIC) as described in Rothan et al. (2019). The first tomato MAGIC population was developed at INRA-Avignon in France and is composed of about 400 lines derived from an 8-way cross (Pascual et al. 2015). This population showed a wide intra-specific genetic variation under control and stress environments and is highly suitable for mapping QTLs (Diouf et al., 2018).

In the present study, we used the 8-way tomato MAGIC population described above and evaluated its response in a multi-environment trial (MET) design. The population was grown in 12 environments including control and several stress conditions (WD, SS and HT), and agronomic traits related to yield, fruit quality, plant growth and phenology were measured. Different plasticity parameters were computed and used together with mean phenotypes to decipher the genetic control of response to environmental variation. Multi-environment QTL analysis was performed in addition to detection of interactive QTLs (QEI) along with QTL mapping for plasticity traits.

## MATERIALS AND METHODS

### Plant material and phenotyping

The MAGIC population was derived from a cross between eight parental lines that belong to *Solanum. lycopersicum* and *Solanum lycopersicum cerasiforme* groups. More details about the population development can be found in Pascual et al. (2015). Briefly, the population was composed of about 400 8-way MAGIC lines that underwent three generations of selfing before greenhouse evaluations were carried out. In this study, a subset of 241 to 397 lines was grown in each environment (Supplemental Table 1).

The full genome of each parental line was re-sequenced and their comparison with the reference tomato genome (‘Heinz 1706’) yielded 4 millions SNPs (Causse et al., 2013). From these polymorphisms, a genetic map of 1345 discriminant SNPs was developed (Pascual et al., 2015) and used in the present study for the QTL analysis.

### Experimental design

The MAGIC population was grown in three different geographical regions (France, Israel and Morocco) and four specific stress treatments were applied. Trials were conducted in order that in a given trial any stress treatment was applied aside a control trial (Supplemental Table 1). Treatments consisted in water deficit (WD), two levels of salinity – considered here as low salinity (LS) and high salinity (HS) – and high temperature (HT) stress. Water deficit was applied by reducing the water irrigation of about 70% and 30% according to the reference evapotranspiration in Israel in 2014 and 2015, respectively and by 50% in Morocco in 2015. Salinity treatment was managed as described in Diouf et al. (2018) and the average electrical conductivity of the substrate (Ec) in Morocco 2016 was 3.76 and 6.50 dS.m^-1^ for LS and HS, respectively; while the Ec in the control condition in Morocco 2015 was about 1.79 dS.m^-1^. For HT stress, plants were sown during the late spring and phenotyped in the summer 2014 in Israel (HIs14) and summer 2017 in France (HAvi17). During HT treatments, greenhouse vent opening was managed all along the entire growing season, with opening the vent only when temperatures rose up to 25°C. Average mean (respectively maximal) temperatures calculated on daily (24 hours) measurements were of 26°C (respectively 34°C) for HAvi17 and 33°C (respectively 48°C) for HIs14. Besides stress treatments, local conventional cultural conditions were applied for control treatments as described in Diouf et al., (2018).

Environments were considered as any combination of a geographical region, a year of trial and an applied treatment (Supplemental Table 1). Climatic sensors were installed in the greenhouses and climatic parameters recorded hourly in all environments. From the climatic parameters, seven environmental covariates were defined (Supplemental Figure 1) including temperature parameters (mean, minimal and maximal daily temperatures and thermal amplitude), the sum of degree-day (SDD), the vapour-pressure-deficit (Vpd in kPa) and the relative humidity (RH) within the greenhouse. To characterize the environments, every covariate was calculated during the period covering flowering time of the population on the fourth truss. Indeed, phenotypic data analyzed here were mostly recorded on the fourth and fifth trusses (Supplemental Table 2). Hierarchical clustering was performed with ‘FactoMineR’ R package (Lê et al., 2008) using the environmental parameters to group environments according to their similarity regarding the within-greenhouse climatic conditions.

**Figure 1:**
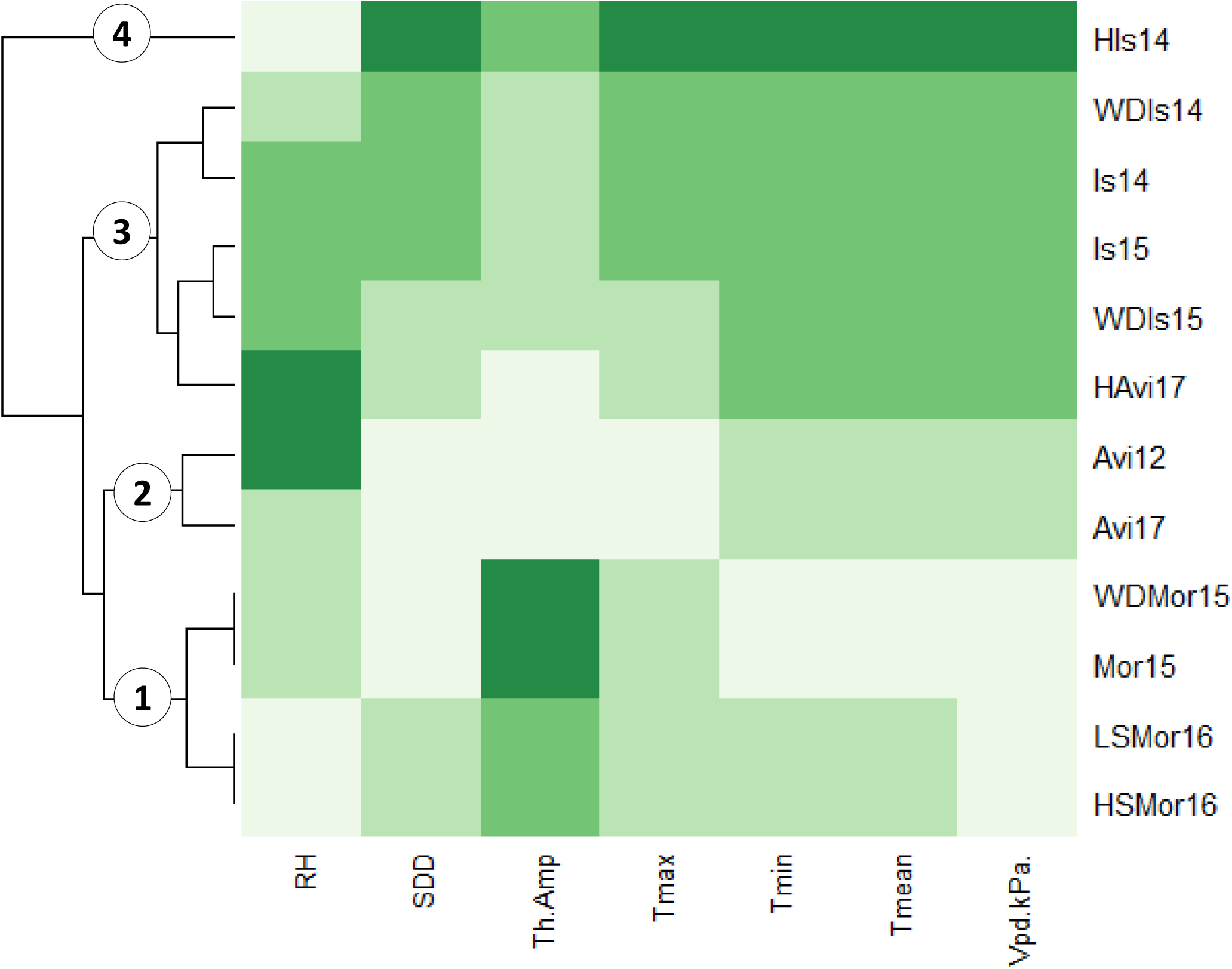
Clustering environments according to seven environmental covariates, measured during the vegetative and flowering stage.

The MAGIC population, the eight parental lines and the four first generation hybrids (one hybrid per two-way cross) were evaluated for fruit weight (FW) by measuring the average FW of the third and/or fourth plant truss in each environment. Phenotypic data were recorded across the different environments for nine supplemental traits related to fruit quality – fruit fruit firmness (firm) and soluble solid content (SSC); plant phenology – flowering time (flw), number of flowers (nflw) and fruit setting (fset); plant development – stem diameter (diam), leaf length (leaf) and plant height (height) and fruit number (nfr). Details about the phenotyping measurements are in Supplemental Table 2. At least two plants per MAGIC line were replicated in each environment except in Avi17 (control condition) where the average phenotype was recorded from single plant measurements. Parents and hybrids had more replicates per genotype (at least two) and served as control lines to measure within-environment heterogeneity.

### Evaluation of GxE and heritability

Data were first analyzed separately in each environment to remove outliers and correct for spatial heterogeneity within the environment. The model (1) below was applied to test for micro-environmental variation within the greenhouse where *y*_*ijjk*_ represents the phenotype of the individual *i*, located in row *j* and position *k* in the greenhouse; *µ* is the overall mean; *C*_*i*_ and *L*_*i*_ represent the fixed effect of control lines and the random effect of the MAGIC lines, respectively. In this model, *t*_*i*_ is an index of 0 or 1, defined to distinguish between control and MAGIC lines; *c*_*ijk*_ is the random residual error.

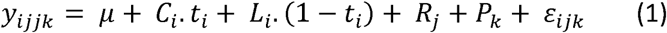

For every trait where row (*R*_*j*_) and/or position (*P*_*k*_) effects were significant, required corrections were applied by removing the BLUP of the significant effects from the raw data. Corrected data were gathered and used in model (2) in order to estimate the broad-sense heritability (*H*^2^) and the proportion of variance associated to the GxE (*prop. σ*^2^_*GxE*_).

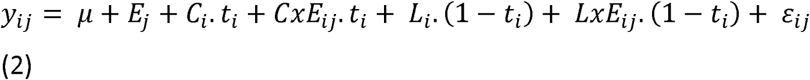

In model (2), *y*_*ij*_ represents the phenotype of the individual *i*, in environment *j*; *µ, C*_*i*_, *L*_*i*_ and the *t*_*i*_ index are as described in model (1); *CxE*_*ij*_ and *LxE*_*ij*_ are the fixed control lines x environment interaction effect and the random MAGIC lines x environment interaction effect, respectively. Within a given environment, random residuals error terms were assumed to be independent and identically distributed with a variance specific to each environment. From this model, the proportion of the total genotypic and GxE variance explained by the model was calculated as the following formula: *prop. σ*^2^_*GxE*_ *= σ*^2^_*LxE*_*/*(*σ*^2^_*L*_ + *σ*^2^_*LxE*_). The significance of GxE was tested with a likelihood ratio test (at 5% level) between the models with and without GxE. The broad-sense heritability at the whole design level (*H*^2^) was derived from variance components of model (2) and calculated as the following: 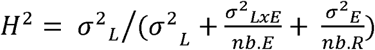, where *σ*^2^_*L*_and *σ*^2^_*LxE*_ are the variance components associated to the MAGIC lines and MAGIC lines x environment interaction effects, respectively. Here *nb. E* and *nb. R* represent the number of environments (*e*.*g*. 12 for FW) and the average number of replicates over the whole design; *σ*^2^_*E*_ is the average environmental variance (*i*.*e*. ∑ *σ*^2^_*Ej*_ */ nb. E*).

### Phenotypic plasticity

Three different parameters of plasticity were estimated using the Finlay-Wilkinson regression (3) and a factorial regression (4) models.

In model (3), *y*_*ij*_ is the phenotype (average values per environment and genotype) and *µ* the general intercept. *G*_*i*_ and *E*_*j*_ are the effects of the MAGIC line *i* and environment *j*, respectively and *β*_*i*_ represents the regression coefficient of the model. It measures individual genotypic sensitivity to the environment.

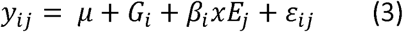

Environments are described here as an index that represents the ‘quality’ of the environment (*i*.*e*. the average performance of all genotypes in a given environment). The *ε*_*ij*_ are the error terms including the GxE and *ε*_*ij*_ ∼ N (0, σ^2^R). From model (3), three parameters were estimated: (i) the genotypic means that is equivalent to the sum (*µ* + *G*_*i*_) representing the average performance of a genotype considering all environments; (ii) he *β*_*i*_ terms (slope), corresponding to genotypic responses to the environments and the variance (VAR) of the *ε*_*ij*_ terms that is a measurement of non-linear plasticity (Kusmec et al., 2017). All these parameters were used then to characterize the genotypes according to their individual performance and their stability in the MAGIC-MET design. For every trait, reaction norms were then computed from the model (3).

The factorial regression model (4) was further applied to describe the GxE through the genotypic response to the different environmental covariates (Tmin°, Tmax°, Tm°, Amp.Th°, Vpd, RH and SDD). The environmental covariates defined from the daily recorded climatic variables in the greenhouses were used for this purpose. For each trait, the most significant environmental covariate (p-value significant at α = 5%) was first identified – by testing successively the significance of each single covariate – and used as an explanatory variable represented by *Cv*_*j*_ in model (4).

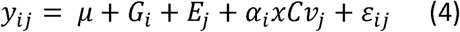

The *α*_*i*_ terms of the model were extracted and considered as a third plasticity parameter (SCv). They represent genotypic sensitivities to the most impacting environmental covariate for each trait. This measurement of plasticity is of interest as it allows identifying the direction and the intensity of each MAGIC line’s sensitivity to a meaningful environmental covariate. Throughout the rest of the document, the ‘slope’ and ‘VAR’ estimated from the Finlay-Wilkinson model and the ‘SCv’ from the factorial regression model will be considered as plasticity phenotypes – all of these parameters being trait-specific.

### Linkage mapping on the genotypic mean and plasticity phenotypes

Linkage mapping was carried out with a set of 1345 SNP markers selected from the genome resequencing of the eight parental lines. All the MAGIC lines were genotyped for those SNPs and at each SNP position, the founder haplotype probability was predicted with the function *calc_genoprob* from R/qtl2 package (Broman et al., 2019). Founder probabilities were then used with the Haley-Knott regression model implemented in R/qtl2 for QTL detection. The response variables were the genotypic means, slope, VAR and SCv for each trait. To attest for significance, the threshold for all phenotypes was set to a LOD threshold of −log10 (α/number of SNPs) where α was fixed at 5% risk level. The VAR plasticity parameter was log transformed for all traits except fset (sqrt transformation) to meet normality assumption before QTL analysis. The function *find_peaks ()* of R/qtl2 package was used to detect all peaks exceeding the defined threshold and the LOD score was dropped of two and one units to separate two significant peaks as distinct QTLs and to define the confidence interval of the QTLs, respectively.

### Multi-environment QTL analysis (QEI)

The strength of QTL dependence on the environment was tested afterward in a second step by identifying QTLs that significantly interact with the environment (QEI). Two multi-environment forward-backward models (5 & 6) were used to test at each marker position the effect of the marker x environment interaction.

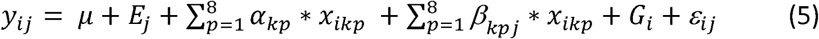

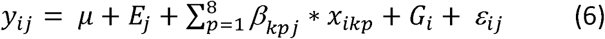

For model (5) and (6), *y*_*ij*_ represents the phenotype (mean value per genotype and per environment), *E*_*j*_ reflects the fixed environment effect; *α*_*kp*_ and *β*_*kpj*_ represent the main and interactive parental allelic effects (*p*)at marker *k* and in environment *j* for *β* _*kpj*_; *x*_*ikp*_ is the probability of the parental allele’s origin for the MAGIC line *i*; *G*_*i*_ stands for a random genotype effect and the residual errors including a part of the GxE that is not explained by the detected QTLs are specific to each environment, *ε*_*ij*_ ∼ N (0, *σ*^2^*Rj*).

Significant QEI were declared in a two-step procedure. First, the main QTL and the QEI effects were tested separately in model (5). The QTL detection process was adapted from the script proposed by Giraud et al., (2017). Every marker showing a significant main QTL or QEI was added as a fixed cofactor and the significance of the remaining markers tested again until no more significant marker was found. All markers selected as cofactors were then jointly tested in the backward procedure and only significant QEI after the backward selection are reported. The second procedure used in model (6) to declare QEI consisted in a slight modification of model (5) where *β* _*kpj*_ represents this time the global (main + interactive) effect of the marker. It allowed the detection of markers that had a main QTL effect or QEI just below the threshold detection but whose global effect is significant when the two components are jointly tested. To determine the threshold level for QEI detection, permutation test were performed 1000 times on the adjusted means with the function *sim*.*sightr* of mpMap 2.0 R package (Huang and George, 2011).

### Data availability

The phenotypic data, average climatic parameters and genotypic information described in the present study are available at https://doi.org/10.15454/UVZTAV. The custom scripts used for the two-stage analysis and QEI modelling are also provided.

## RESULTS

### Environment description

The 12 environmental conditions were described by the daily climatic parameters recorded until the end of flowering of the 4^th^ truss. Seven environmental covariates were selected, and the environments clustered according to these covariates in four groups (Figure 1). The first group included all trials from Morocco that were characterized by high thermal amplitude and low Vpd. The control environments in France (Avi12 and Avi17) clustered together in the 2^nd^ group, defined by low maximal temperatures and high relative humidity. HIs14 clustered alone in the 4^th^ group and formed the most extreme environment showing very high temperatures and dry climate with low relative humidity. The remaining environments clustered together in the 3^rd^ and most disparate group.

Phenotypic distributions were plotted for each trait regarding the environments where it was evaluated (Supplemental Figure 2) showing a distribution in accordance with the clustering of the environments for some traits (firm, height, nflw and leaf). Other traits such as FW, nfr, SSC and fset showed a distribution pattern with relatively high within-group variability, notably for environments clustering in group 1 from Morocco.

**Figure 2:**
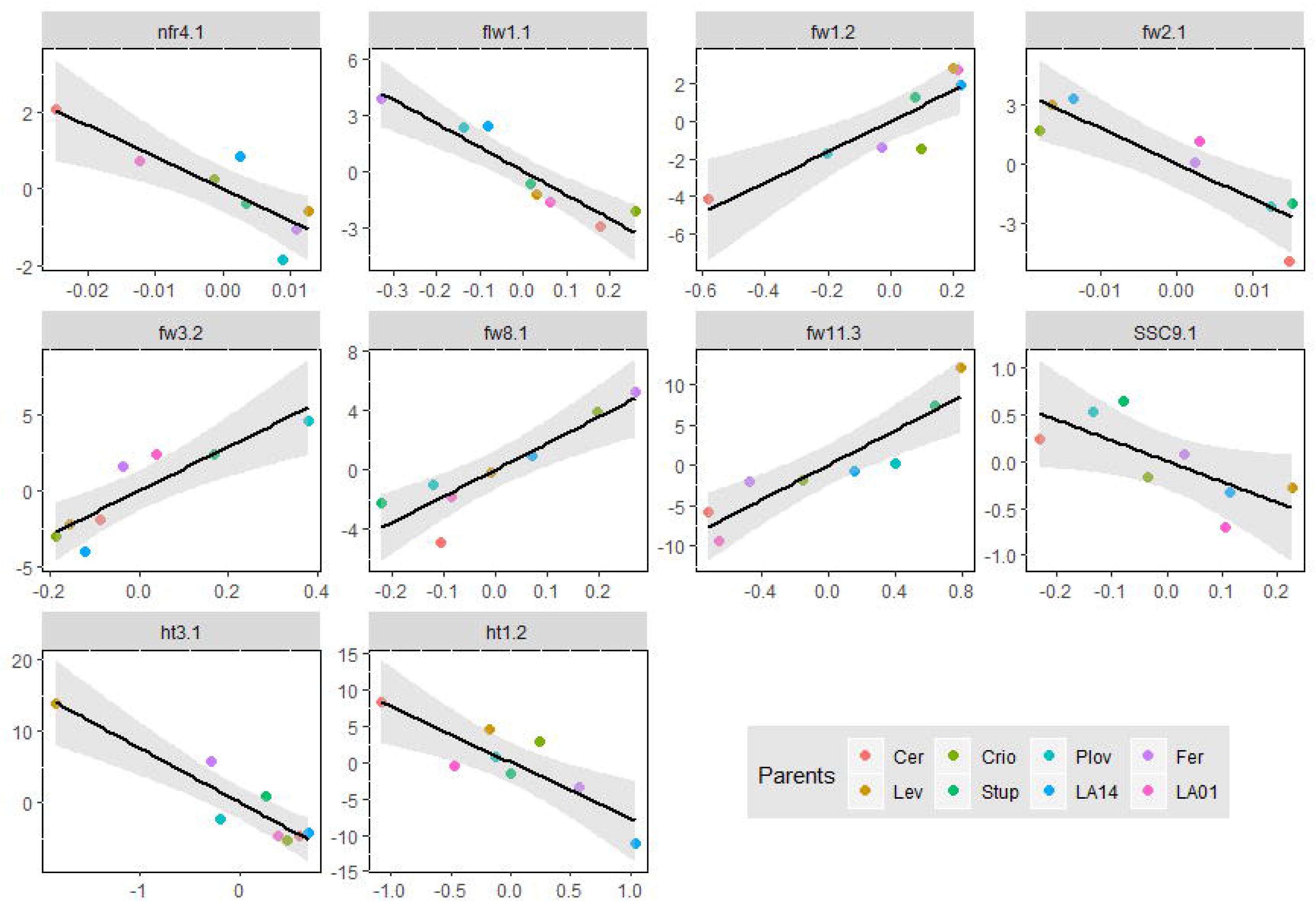
Pearson’s correlation between mean and plasticity parameters.

### GxE in the MAGIC population

Genotype x environment interaction analysis was carried out after correcting data for micro-environmental heterogeneity and removing outliers. As a first step, variance analysis was conducted with ASReml-R package and the variance components from model (2) used to estimate the proportion of GxE variance (*prop. σ*^2^_*GxE*_) and heritability at the whole design level (*H*^2^). Significant GxE was found for every trait and the *prop. σ*^2^_*GxE*_ varied from 0.15 (for nflw) to 0.68 (for leaf). Although GxE was significant, seven out of the ten measured traits showed a higher proportion of genotypic variance compared to GxE (Supplemental Table 3). The broad-sense heritability of the whole design *H*^2^ was largely variable according to the trait, varying from 0.18 (nfr) to 0.77 (flw). Its calculation took into account the residual environment-specific variance which showed different range according to the trait, lowering heritability of traits such as nfr and fset (Supplemental Table 3). Furthermore, *H*^2^ at the whole design level was lower than the heritability computed in single environment (Supplemental Figure 3).

**Figure 3:**
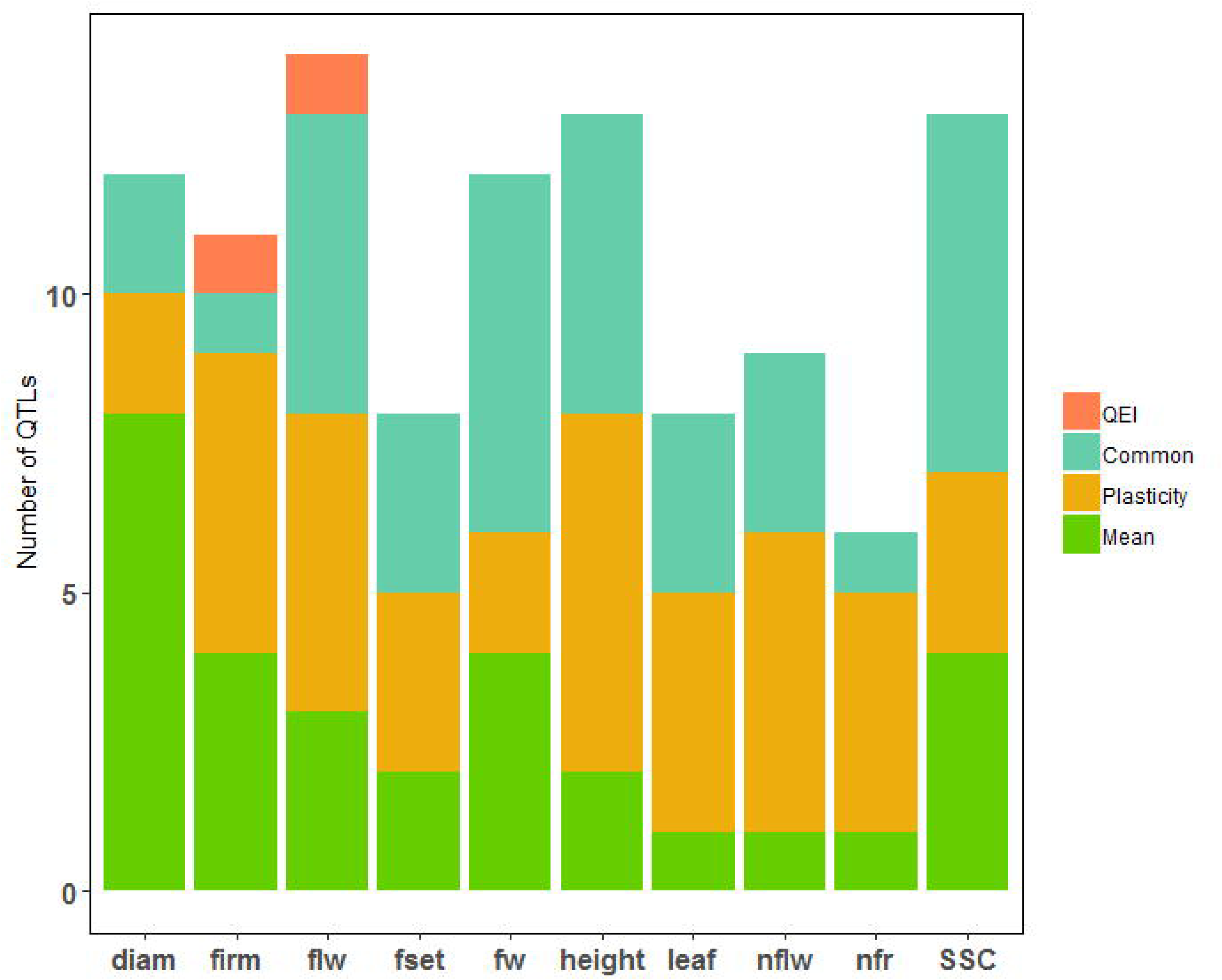
Representation of plasticity QTLs along the genome. Numbers above the square represent the different chromosomes and the colors distinguished the different traits. The x-axis represents the physical distances in mega base pair (Mbp).

Afterwards, the proportion of the GxE that could be predicted by the environmental covariates was assessed following the factorial regression model (4). Across traits, different environmental covariates significantly explained the GxE #(Supplemental Figure 4). Considering only the most significant covariate, from 18% (FW) to 47% (fset) of the GxE (proportion of the sum of squares) could be reliably attributed to the responses of genotypes to climatic parameters measured within the greenhouses. To perform the factorial regression model (4), the most important environmental covariate was first identified for each trait (Supplemental Figure 4). Growth traits, height and leaf were for example mostly affected by the thermal amplitude and maximal temperature, respectively, while yield component traits, FW and nfr were particularly sensitive to the sum of degree day. The vapour pressure deficit (Vpd, kPa) was the most important environmental factor affecting firm, fset and SSC. Flowering time (flw) and nflw were mostly affected by minimal temperatures and relative humidity, respectively. Stem diameter was the only trait for which none of the environmental covariates significantly affected the trait.

**Figure 4:**
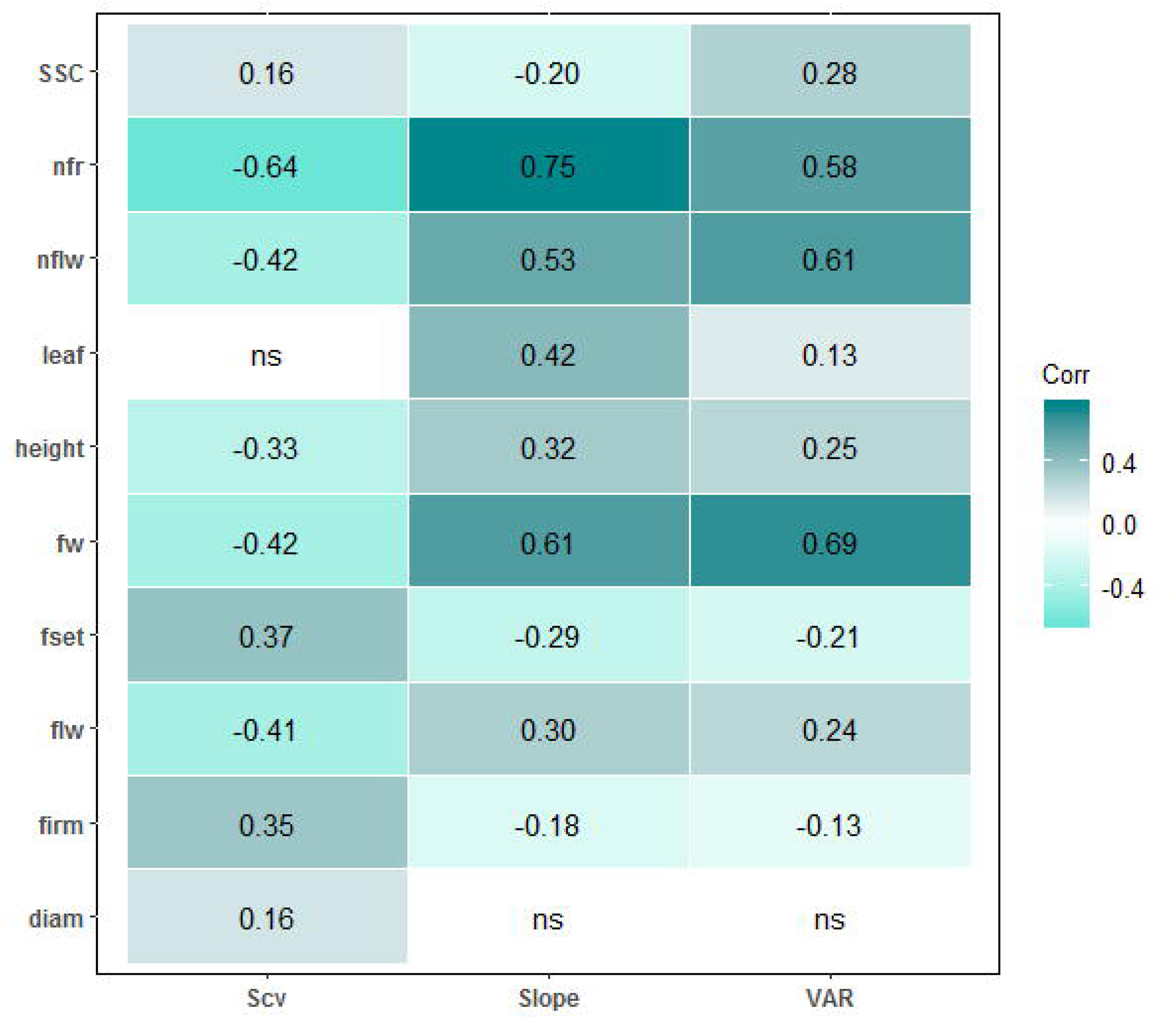
Number of QTLs identified specifically on mean, plasticity or QEI and QTLs that were common to at least two of them.

### Phenotypic plasticity

Three different parameters were used to quantify phenotypic plasticity in the MAGIC-MET design. For each trait, the slope and VAR from the Finlay-Wilkinson regression model and the genotypic sensitivity to the most important environmental covariate (SCv) from the factorial regression model were extracted. A large genetic variability was observed for plasticity of all traits (Supplemental Figure 5 and Supplemental Figure 6). Besides, significant correlations were found between the mean phenotypes and plasticity parameters (Figure 2) for most of the traits. The best average-performing genotypes were usually the most responsive to environmental variation as highlighted by the positive correlation between the genotypic means and slope from the Finlay-Wilkinson regression model. The majority of the MAGIC lines responded in the same direction to the environmental quality and only a few genotypes (none in the case of height) showed negative reaction norms; however, more divergent shapes of reaction norms were observed from the factorial regression model (Supplemental Figure 5).

**Figure 5:**
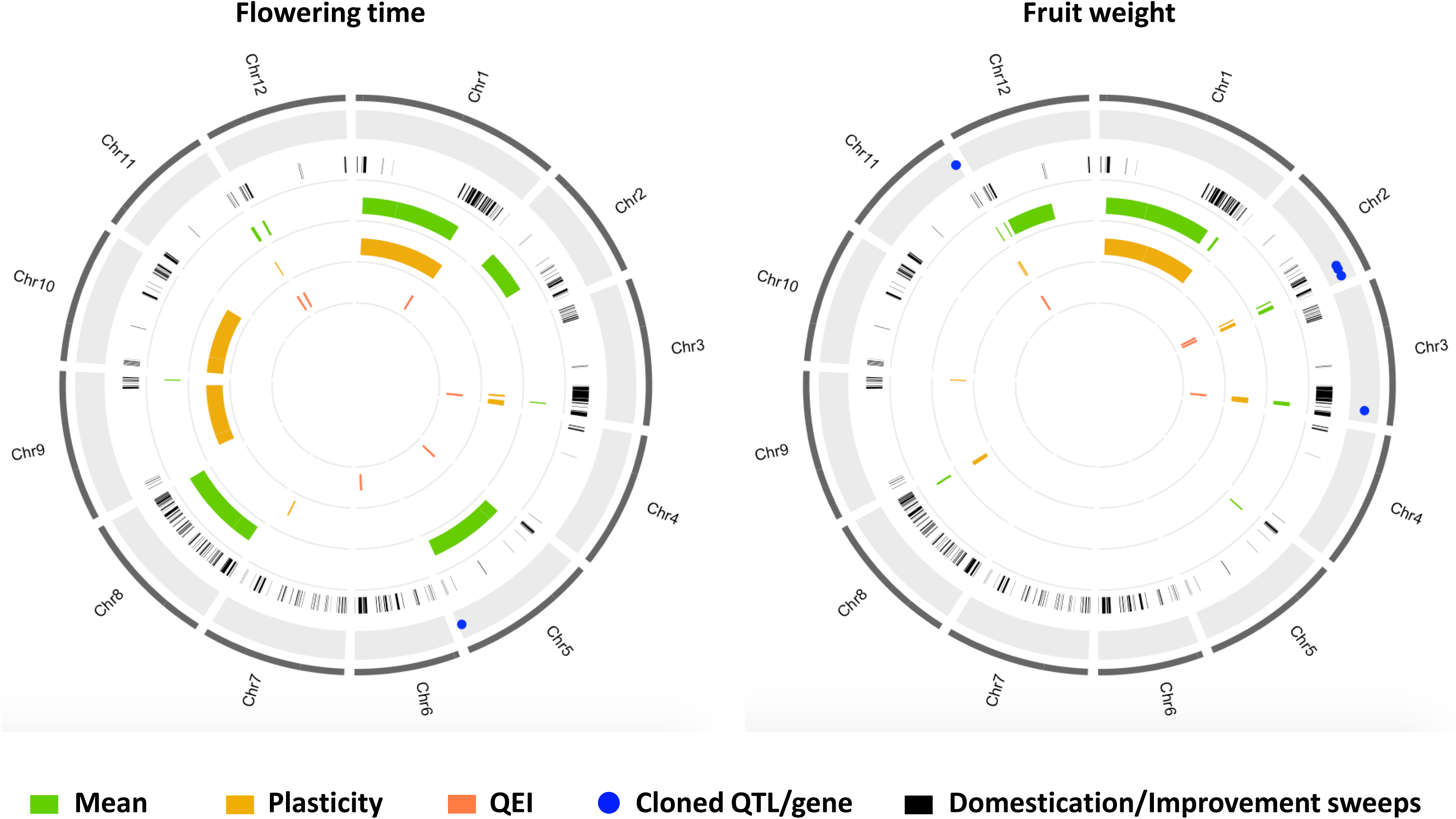
Physical positions of the MAGIC-MET QTLs for fruit weight and flowering time. The following circle with black bars represents the different domestication/improvement sweep regions identified in (Zhu et al. 2018). The other circles plot the CI of QTLs identified on mean (green), plasticity (orange) or with QEI analysis (purple).

**Figure 6:**
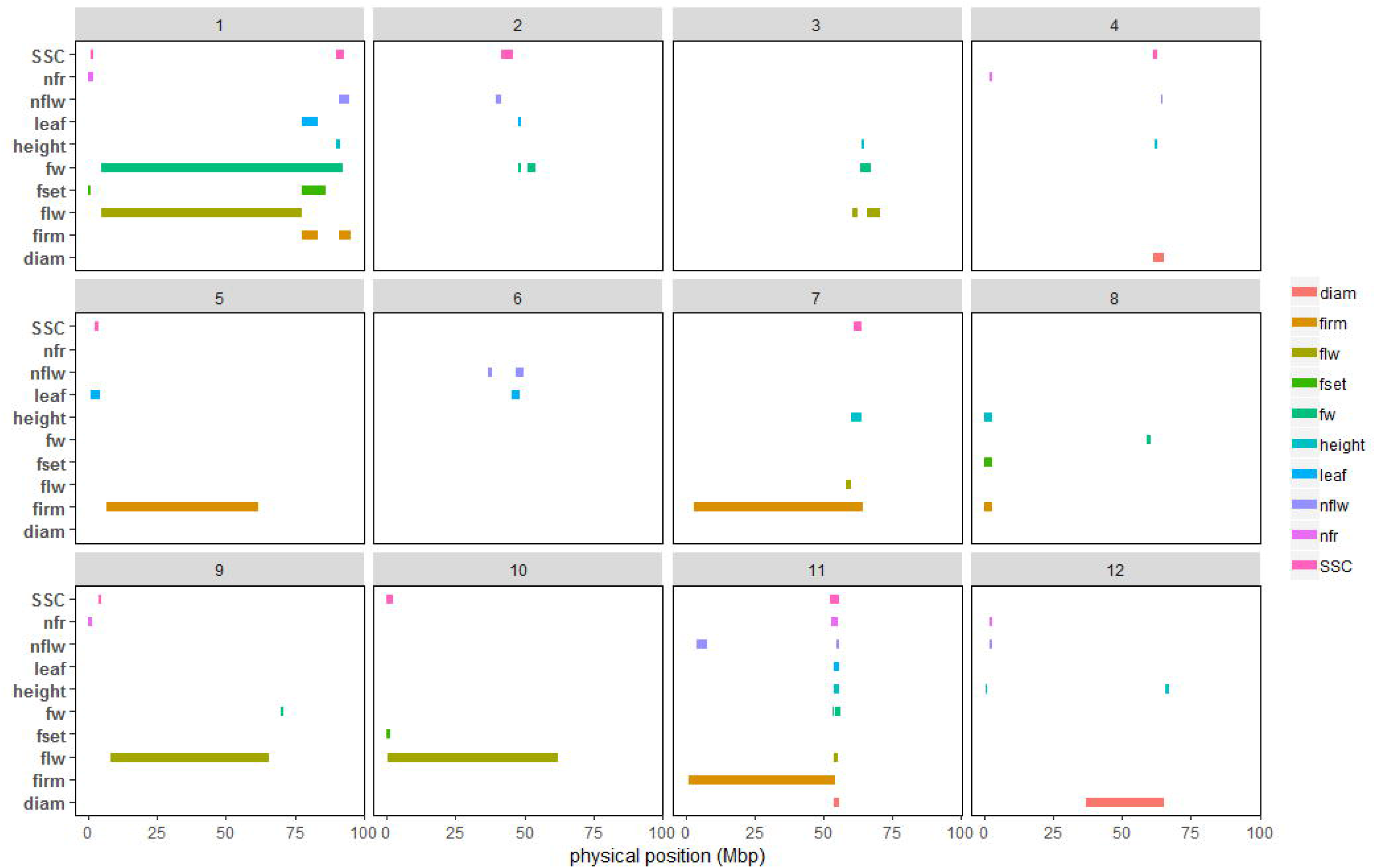
Correlation of the estimated allelic effect for consistent QTLs between mean and plasticity phenotypes.

### QTL mapping

We used genotypic means and plasticity measurements for every trait as input phenotypes to decipher the genetic architecture of tomato response to abiotic stresses. Considering the 10 traits evaluated, a total of 104 unique QTLs were identified for genotypic means and the plasticity parameters (Supplemental Table 4). The proportion of QTLs shared between mean and plasticity was about 21%, lower than QTLs that were plasticity or mean specific (79%). Considering only the 63 plasticity QTLs, 11 and 7 QTLs were specifically detected with the SCv and VAR plasticity parameters. Plasticity QTLs were detected on every chromosome (Figure 3); however, the chromosome 1 showed the highest number with 12 plasticity QTLs. In this chromosome, plasticity QTLs were detected at least once for every trait. The chromosome 11 carried a total of 11 plasticity QTLs and interestingly all these QTL (except ppnflw11.1) co-localized in a short region of the chromosome between 52 and 55 Mbp. The chromosomes 5, 6 and 10 showed the lowest number (only 3) of plasticity QTLs. For QTLs detected on genotypic means, the number of QTLs per chromosome varied from 2 QTLs on chromosomes 6 and 10 to one QTL on chromosome 1.

### QTL-by-environment analysis (QEI)

Multi-environment forward-backward models were used to assess the significance and the strength of the QTL effects across environments. The QEI analysis was conducted in two steps using the same set of 1345 SNP markers that were also used for linkage mapping analysis. This analysis yielded 28 QEI (only those showing significant interaction) for the 10 traits (Supplemental Table 4). The number of QEI varied from 0 QEI for nfr to 6 QEI for flw. These two traits also demonstrated the lowest and highest *H*^2^.

All QEI identified in this step were confronted to the plasticity and genotypic means QTLs using the physical positions of the QTLs and their confidence intervals. Interestingly, this comparison revealed that all the detected QEI were also identified using either genotypic means or plasticity parameters, in the linkage mapping analysis, except two QEI located on the same region of chromosome 6 (flw6.1 and firm6.1). Among the 106 unique QTLs identified on genotypic means, PP and QEI, a notable number of QTLs were specific representing 30 and 32% for plasticity and genotypic means, respectively (Figure 4). Eight QTLs involving five different traits (flw1.1, fw2.1, fw2.2, fw11.2, leaf6.1, nflw11.2, SSC1.2 and SSC9.1) were identified with all the three approaches highlighting their robustness and susceptibility to environmental variation.

### Genetic location of the MAGIC-MET QTLs

The physical positions based on the SL2.50 version of the reference genome, were used to compare the position of the different QTL category (genotypic means, plasticity or QEI). Indeed, a recent study has identified different tomato regions (Sweep regions) that were selected during domestication and improvement events (Zhu et al., 2018). These regions were cross checked against the positions of our QTLs. Some QTLs detected in the MAGIC-MET design were located in large regions thus colocating with a high number of Sweep regions (Figure 5 & Supplemental Figure 7). Thus, considering only the QTLs with CI lower than 2Mbp intervals and all QEI, a total of 61 QTLs were selected and compared with the Sweep regions. Plasticity QTLs appeared to be in majority located within the Sweep regions and only 6% of the selected plasticity QTLs were outside the domestication/improvement selective sweeps (Supplemental Figure 8). Interestingly, the Sweep region SW75 located in chromosome 3 (between 64.76 and 65.01 Mbp) carried a total of five QTLs (ht3.1, fset3.1, flw3.2, leaf3.1, fset3.1). The Supplemental Table 5 presents all the Sweep regions holding at least one MAGIC-MET QTL. Chromosome 11 was highlighted as holding a number of plasticity QTLs for different traits (Figure 3). Indeed, seven different QTLs all identified with plasticity parameters, were located within the Sweep regions SW254 and SW255, from 53.81 – 55.62 Mbp on chromosome 11 (Supplemental Figure 9). Among the ten QTLs that were outside the Sweep regions, one QTL was identified for mean FW and located on chromosome 5 (fw5.1) in position 4.52 Mbp. This QTL was mapped in a region holding other QTLs segregating in the MAGIC population for fruit size, fruit width and fruit length (Supplemental Table 6; data from the experiment in Pascual et al. (2015)).

### Candidate genes

Confidence intervals (CI) of the MAGIC-MET QTLs varied from 0.45Mbp to 87Mbp including a variable number of genes. We thus focused on QTLs presenting CI regions smaller than 2Mbp for CG screening. From 49 (nflw12.1) to 256 (diam4.1) genes were within the regions of the selected QTLs. Taking advantage of the parental allelic effect, the CG were narrowed for each QTL by contrasting the allelic effect of the eight parental lines. The selected candidates after the filtering procedure are presented in Supplemental Table 7, highlighting interesting candidates for further studies. Flowering time QTLs for instance included some CG with consistent matching regarding their functional annotation. For example, the CI of the QTL ppflw11.1 on chromosome 11 included two CG: Solyc11g070100 and Solyc11g071250 corresponding to “Early flowering protein” (ELF) and “EMBYO FLOWERING 1-like protein” (EMF1), respectively. Among other potential flowering candidates, we noticed Solyc12g010490 (AP2-like ERF) for the QTL flw12.1 and Solyc03g114890 and Solyc03g114900 (COBRA-like proteins) for the QTL flw3.2. Aside flowering time, the selected candidate genes for the QTLs diam4.1 and ppSSC1.1 included the Solyc04g081870 (annotated as an Expansin gene) and Solyc01g006740 (annotated as Sucrose phosphate phosphatase) genes, respectively.

We could identify some plasticity QTLs showing sensitivity to the environmental conditions, notably the QTLs detected using the Scv plasticity parameter. Candidate genes were screened for some QTLs falling into this category. The ppfw9.1 QTL CI for example, showing susceptibility to the sum of degree day (SDD), carried a chaperone candidate (solyc09g091180) which might be involved in regulating fruit weight depending on the SDD variation. Similarly, the QTL ppleaf11.1 is affected by the maximal temperature (Supplemental Table 4). Three CG (Solyc11g071830, Solyc11g071930 and Solyc11g071710) belonging to the Chaperone J-domain family, were retained after the filtering procedure in the region of this QTL. Interestingly, the DnaJ-like zinc-finger gene (Solyc11g071710) was among the candidates corresponding to several plasticity QTL including ppflw11.1, ppleaf11.1, ppnflw11.1, ppht11.1 and ppdiam11.2. This gene presented a total of 122 polymorphisms across the eight parental lines among which 35 and 68 are in the up-stream and down-stream gene region. Further investigation regarding this gene is needed to state its potential pleiotropic effect.

## DISCUSSION

### Genetic variability in tomato response to environmental variation

Genotype x environment interaction is a long-standing challenge for breeders and the predicted climate change has encouraged plant geneticists to devote more attention into understanding its genetic basis. Tomato is a widely cultivated crop adapted to a variety of environmental conditions (Rothan et al. 2019). However, important incidences of abiotic stress in the final productivity, fruit quality and reproductive performance have been noticed (Albert et al. 2016; Estañ et al. 2009; Mitchell et al. 1991; Xu et al. 2017). We quantified the level of GxE and the subjacent phenotypic plasticity in a multi-environment and multi-stress trial – involving induced water-deficit, salinity and heat stresses – in a highly recombinant tomato population. An important genetic variability was observed for the plasticity traits related to yield, fruit quality, plant growth and phenology (Supplemental Figure 6). This highlights the interest of the MAGIC population as a valuable resource for tomato breeding in dynamic changing environments. Tomato wild species have been also characterized as an important reservoir for abiotic stress tolerance genes (Foolad, 2007). However, their effective use in breeding programs could be difficult due to undesirable linkage drag, notably for fruit quality. Unlikely, the MAGIC population characterized here is an intra-specific population with high diversity regarding fruit quality components, which provides a great advantage as a breeding resource compared to wild populations.

Several statistical models are available to explore, describe and predict GxE in plants (Yan et al., 2007; Malosetti et al., 2013). Factorial regression model is among the most attractive as it allows to describe the observed GxE regarding relevant environmental information. We used the factorial regression model with different environmental covariates that are readily accessible from year to year, which allowed us to predict a variable proportion of the observed GxE (Supplemental Figure 4). Besides, each MAGIC line was characterized for its sensitivity to the growing climatic conditions opening avenues to effectively select the most interesting genotypes for further evaluation in breeding programs targeting stressful environments.

Interestingly we found significant correlation between the genotypic sensitivities to the different environmental covariates and slopes from the Finlay-Wilkinson regression model (Supplemental Figure 10). This emphasizes the adequacy of the selected environmental covariates to explain differences observed in the average performance of the genotypes across environments. Conversely, slope and VAR showed less significant correlations, although they were both correlated to mean phenotypes in the same direction – except for SSC (Figure 2). This may be induced by distinct genetic regulation of these two plasticity parameters which reflect different types of agronomic stability (Lin et al. 1986). Indeed, we identified 7 and 14 plasticity QTLs that were specific to VAR and slope, respectively (Supplemental Table 4). The correlation pattern of the different plasticity parameters evokes a complex regulation of plasticity which besides is seemingly trait specific.

Significant correlation at phenotypic level might result from the action of pleiotropic genes. The Figure 2 displays the correlations between genotypic means and plasticity which were significant for almost every trait at variable degree. These correlations were reflected at the genetic level by 22 QTLs overlapping between genotypic mean and plasticity parameters, representing about 21% of all identified QTLs. However, a high proportion of the QTLs were specific either to genotypic means or plasticity parameters (Supplemental Figure 11), hence suggesting the action of both common and distinct genetic loci in the control of mean phenotype and plasticity variation in tomato.

### Genomic location of the MAGIC-MET QTLs

The availability of substantial genomic information in tomato enabled the identification of different genomic regions which have undergone selective sweeps which were strongly selected during the domestication and improvement process (Lin et al. 2014; Zhu et al. 2018). When projected on the physical positions of the tomato reference genome (SL2.50 version), most of the plasticity QTLs we identified were located within the sweep regions defined by Zhu et al. (2018). It therefore suggests that plasticity might have been selected together with other interesting agronomic traits during tomato domestication and improvement. For instance, this is corroborated by the positive correlation between slope (from the Finlay-Wilkinson regression model) and mean fruit weight variation. Indeed, genotypes with higher FW slope are characterized by good adaptability in high quality environments and will likely be intended to selection. Co-selection of allelic variants leading to higher performance in optimal condition together with plasticity alleles is a realistic assumption that would explain the significant correlation that we observed between the genotypic means and plasticity. In rice for instance, *GhD7* has been described as a key high-yield gene simultaneously involved in the regulation of plasticity of panicle and tiller branching and involved in abiotic stress response (Herath 2019). This example highlights a gene carrying different allelic variants affecting together plasticity and mean phenotypes. Further investigations are needed to assess how domestication and breeding have affected plasticity in tomato and other crop species.

An important genomic region involved in the genetic regulation of plasticity for six different traits was identified in chromosome 11 (Supplemental Figure 9). This region is obviously a regulatory hub carrying interesting plasticity genes. It remains to determine if the co-localization of the different plasticity QTLs in this region is due to the action of a pleiotropic gene or different linked genes. Nevertheless, the chromosome 11 region highlighted here is an interesting target for breeding as well as for understanding the functional mechanisms of plasticity genes.

### Allelic-sensitivity vs gene-regulatory model

Sixty-three plasticity QTLs were identified among which 22 (35%) were also identified when using the genotypic means; and 41 (65%) were specific to plasticity. Via et al. (1995) proposed two genetic models – the allelic-sensitivity and gene-regulatory models – among the mechanisms involved in the genetic control of phenotypic plasticity. These two models are distinguishable through QTL analysis (Ungerer et al., 2003) with the expectation that allelic-sensitivity model will lead to co-localization of genotypic means and plasticity QTLs, while a distinct location of QTLs affecting mean and plasticity will likely correspond to the gene-regulatory model (Kusmec et al., 2017). Regarding our results, both models are suspected to regulate tomato plasticity, even though the gene-regulatory model is predominant with 65% of the plasticity QTLs that did not co-localize with genotypic means QTLs for the same trait. In maize, using a larger number of environments and traits, Kusmec et al. (2017) found similar results and even a higher rate of distinct locations of plasticity and mean QTLs. Studying plasticity as a trait *per se* is therefore of a major interest since breeding in both direction (considering the mean phenotype and its plasticity) is achievable. Through transcriptomic analyses, Albert et al. (2018) observed that genotype x water deficit interaction was mostly associated to *trans-*acting genes which could be assimilated to the gene-regulatory model in agreement with our results.

Although the distinct location of QTLs detected on plasticity and genotypic means could be confidently assigned to the action of genes in interaction, their co-localization is not necessarily a case of allelic-sensitivity regulation, especially if the QTL is in a large region. Indeed, the allelic-sensitivity model assumes that a constitutive gene is directly sensitive to the environment regulating its expression across different environmental conditions, inducing hence phenotypic plasticity. This is a very strong hypothesis regarding the QTLs since the overlapping region between genotypic means and plasticity could carry different causal variants in strong linkage disequilibrium affecting either mean phenotype or plasticity. Thus, co-locating mean and plasticity QTLs should be not automatically imputed to the allelic-sensitivity model. We found a total of 22 constitutive QTLs between genotypic means and plasticity for all 10 measured traits (Supplemental Table 4). Considering the estimated QTL effects, the variation patterns of the eight parental allelic classes were compared between mean and PP QTL of the same trait. Only ten QTL showed consistent allelic effects (Spearman correlation significant at 0.05 threshold level) strengthening the hypothesis of the allelic-sensitivity model for these QTLs (Figure 6). Further studies should help to elucidate and validate the candidate plasticity genes and to clarify their functional mechanism.

### Complementary methods to identify environment-responsive QTLs

Different approaches have been proposed in the literature to dissect GxE into its genetic components (Malosetti et al., 2013; El-Soda et al., 2014). We used a mixed linear model with a random genetic effect accounting for the correlation structure of the MAGIC-MET design to identify the QEI. Extending the use of mixed linear models to MAGIC populations in the framework of MET analysis has been very rarely applied in crops. To our knowledge, only Verbyla et al., (2014) applied such approach in wheat and identified QEI for flowering time. Our model was adequate to account for the complex mating design of the MAGIC population by using the haplotype probabilities. Indeed, it allows estimating the QTL effect for each parental allelic class and for each environment at every SNP marker. Overall, 28 QEI were detected showing significant marker x environment interaction for ten traits.

Methods using plasticity as a trait *per se* are also attractive to identify environmentally sensitive QTLs. This strategy was applied in maize, sunflower, barley and soybean to detect the loci governing GxE (Lacaze et al., 2009; Gage et al., 2017; Kusmec et al., 2017; Mangin et al., 2017; Xavier et al., 2018). With different plasticity parameters, we identified a total of 63 plasticity QTLs and only 24% were also identified with the QEI models. Thus, both methods, using plasticity or mixed linear models, are complementary approaches to study the genetic component of GxE.

### Candidate genes

Multi-parental populations are powerful for QTL mapping studies (Huang et al. 2012; Kover et al. 2009) and are besides interesting for fine mapping and candidate gene screening. Barrero et al. (2015) for instance considered the variation of the QTL effect estimated for the different parental lines, combined with transcriptomic analyses to efficiently identify candidate genes. Similarly, Septiani et al. (2019) narrowed candidate genes for Fusarium resistance in a maize MAGIC population using allelic effect of the MAGIC parents.

A number of candidate genes were proposed in our study, affecting both genotypic means and plasticity variation. These candidate genes were selected based on the parental allelic effect and represent valuable targets for future studies attempting to characterize the molecular mechanisms underlying plasticity in tomato. Indeed, relevant candidate genes were proposed for plasticity of flowering time including the Solyc11g071250 which corresponds to an “EMBYO FLOWERING 1-like protein” (EMF1). The implication of EMF1 in flowering time has been observed in Arabidopsis by Aubert et al., (2001) who highlighted an indirect effect of EMF1 on flowering time and inflorescence. More recently, Luo et al., (2018) outlined the role of EMF1 interacting with CONSTANS proteins in a complex pathway to regulate the expression of flowering time genes in Arabidopsis. Solyc11g070100 which is annotated as “Early flowering protein” (ELF) gene is also an interesting candidate for flowering time regulation. It was observed across species that a consistent expression of ELF3 can extend the rapid transition to flowering (Huang et al., 2017). ELF3 loss of function is therefore expected to trigger early flowering according to these authors. Interestingly, Solyc11g070100 was affected by 69 SNPs and 14 INDELs polymorphisms, among which only one SNP showed polymorphism variation in line with the estimated allelic effect for the eight parental lines at this QTL. This SNP was localized at the position 54,632,225 bp in chromosome 11, upstream the gene Solyc11g070100. The parent LA1420 carried the reference allele at this SNP while the remaining parents held the alternative allele. Considering the estimated allelic effects at this QTL, we could assume that the LA1420 allele variant might induce an early flowering phenotype comparatively to the other parents.

### Conclusion

We aimed to dissect the genetic architecture of tomato response to different environments involving control and stress growing conditions. The MAGIC population demonstrated a large genetic variability in response to abiotic stresses which was reflected by the identification of 63 plasticity QTLs. This was achieved through the use of different plasticity parameter highlighting the importance of plasticity quantification for deciphering its genetic basis. The plasticity QTLs were in majority (65% of the plasticity QTLs) located in distinct regions than the QTLs detected for the mean phenotypes, suggesting a specific genetic control of mean trait variation and plasticity at some extent. Using plasticity as a trait *per se* in mapping analysis turned out to be a good method for identifying genetic regions underlying GxE. Almost all the QEI were also identified for at least one of the plasticity parameters. Overall, this study presents the MAGIC population as a powerful resource for tomato breeding under abiotic stress conditions, as well as for understanding the genetic mechanisms regulating tomato response to environmental variation.

## Acknowledgements

We acknowledge the greenhouse staff of INRA GAFL, Gautier SEMENCES and Hazera seeds for the trial management. The ANR project Adaptom (ANR-13-ADAP-0013) and TomEpiSet (ANR-16-CE20-0014) supported this work. ID was supported by the WAAPP (West Africa Agricultural Productivity Project) fellowship and hosted as Ph.D. student in the INRA GAFL.

## Supplemental files

**Supplemental Figure 1:** Selection of 7 environmental covariates for the factorial regression model. Three periods – each of 20 days – were defined from planting to the end of flowering on the 4^th^ truss. The period from 20 to 60 days after planting (DAP) covered vegetative growth and flowering on the 4^th^ truss and the measured climatic variables averaged during this period. The different environmental covariates are described

**Supplemental Figure 2:** Boxplot distribution of the traits across environments. The colors of the boxplot are according to the groups defined by clustering of the environments

**Supplemental Figure 3:** Heritability in the MAGIC-MET design. For each trait, heritability was computed at every environment and plotted with heritability of the full design ***H***^**2**^ (in green)

**Supplemental Figure 4:** Proportion of the sum of square attributed to the different factors in the factorial regression model. For each trait, the orange and green stacked bars represent the proportion of the SSq explained by the Genotype and Environment factors in model (4). The remaining colors represent the effect part of the GxE that could be explained by the different environmental covariates. Only significant covariates were highlighted within the bars.

**Supplemental Figure 5:** Reaction norms from the Finlay-Wilkinson regression model (A) and the factorial regression model (B). In figure 5 A, the blue and orange lines represent the positive and negative reaction norms. In Figure 5 B, the green and purple lines represent the positive and negative reaction norms

**Supplemental Figure 6:** Histogram distribution of mean and all plasticity parameters for each trait

**Supplemental Figure 7:** Physical positions of the MAGIC-MET QTLs for diam, leaf, height, fset, nflw, nfr, firm and SSC. The outer circle with gray font represents the known and cloned QTL/gene for each trait. The following circle with black bars represents the different domestication/improvement sweep regions identified in (Zhu et al. 2018). The other circles plot the CI of QTLs identified on mean (green), plasticity (orange) or with QEI analysis (purple)

**Supplemental Figure 8:** Number of the MAGIC-MET QTLs identified within or outside the domesticated/improved regions. Only the MAGIC-MET QTLs within short CI (lower than 2Mbp) were considered. The response specific category included QEI and plasticity specific QTLs; the common category correspond to QTLs that were commonly identified on mean, plasticity and QEI or at least two of them

**Supplemental Figure 9:** Zoom plot on Chromosome 11 region from 53-57 Mbp. Each color represents a different QTL located in this region and the top black bars are the Sweep regions SW254 and SW255

**Supplemental Figure 10:** Correlation between the genotypic sensitivities to environmental covariates from the factorial regression model and slopes from the Finlay-Wilkinson regression model

**Supplemental Figure 11:** Venn diagram of the number of QTL specific or commonly detected with mean, PP or using the QEI models.

**Supplemental Table 1:** Description of the MAGIC-MET design with the 12 environments and their respective names

**Supplemental Table 2:** Description of the phenotypic traits evaluated in the MAGIC-MET design

**Supplemental Table 3:** Estimates of the variance components from model (2)

**Supplemental Table 4:** Results of QTL and QEI analysis in the MAGIC-MET design

**Supplemental Table 5:** Genetic location of the MAGIC-MET QTLs overlapping with the Sweep (domestication/improvement) regions.

**Supplemental Table 6:** QTLs identified for fruit size, fruit width and fruit length in the MAGIC population

**Supplemental Table 7:** Selected candidate genes for all the mean and plasticity QTLs located within 2Mbp CI region

